# The pharmacopeia of helminth-derived metabolites driving tolerance in dendritic cells

**DOI:** 10.1101/2023.01.26.525718

**Authors:** Aubrey N. Michi, Nathalia L. Malacco, Jose A. Sandoval, Elizabeth Siciliani, Ana G. Madrigal, Tamara Sternlieb, Ghislaine Fontes, Irah L. King, Igor Cestari, Armando Jardim, Mary M. Stevenson, Tony Hunter, Fernando Lopes

## Abstract

Helminth infections can restore immune tolerance and suppress intestinal inflammation, yet the molecular mediators underlying these effects are poorly defined. Here, we show that nonpolar metabolites derived from the helminth *Heligmosomoides polygyrus bakeri* (HnpM) program dendritic cells (DCs) toward a tolerogenic DC (tolDC) phenotype. Adoptive transfer of HnpM-treated DCs significantly ameliorated DSS-induced intestinal inflammation in mice. Transcriptomic and metabolomic profiling revealed that HnpM induces early metabolic reprogramming followed by gene expression signatures characteristic of tolerogenic DCs that are shared by established tolDC-inducing agents. Together, these findings identify that helminth-derived metabolites are a novel source of small molecule therapeutics to restore immune tolerance in immune-mediated disorders.

**Summary Sentence:** Metabolites produced by *Heligmosomoides polygyrus* induce metabolic and transcriptional changes in DCs consistent with tolDCs, and adoptive transfer of these DCs attenuate DSS-induced intestinal inflammation.

## INTRODUCTION

Inflammatory bowel diseases (IBD) arise in genetically susceptible individuals in which environmental triggers disrupt intestinal immune tolerance, leading to exaggerated CD4^+^ Th1 and Th17 immune responses(1). Helminth infections, including *Heligmosomoides polygyrus bakeri* (Hpb), are known to restore immune tolerance and suppress intestinal inflammation by modulating CD4^+^ T cell immunity(2). It has been shown that they induce IL-10 and TGF-β–producing lamina propria T cells, which confer protection to colitis by mesenteric lymph node T cells, and reduce Th1/Th17-driven pathology in Hpb-infected mice.(2) As dendritic cells (DCs) shape CD4^+^ T cell polarization(3), many of the immunomodulatory effects of helminths are mediated through DC reprogramming, with helminth-exposed DCs acquiring tolerogenic phenotypes that limit inflammatory T cell responses(4). Here, we examined whether helminth-released metabolites directly promote immune tolerance, thereby opening new avenues to explore these small molecules as a pharmacopeia for the treatment of inflammatory diseases.

## RESULTS & DISCUSSION

To test whether helminth-derived metabolites induce tolerogenic DCs (tolDCs), we biochemically fractionated and isolated polar (HpM) and nonpolar (HnpM) metabolites from Hpb culture supernatants and treated B6 mouse bone marrow–derived DCs (BMDCs) with each fraction for 20h, followed by a 4h LPS challenge **(Fig. 1A)**. Whereas LPS alone induced robust TNF production, HnpM, but not HpM or mock extracted fractions, significantly reduced TNF and enhanced IL-10 release **(Fig. 1B, C)**, with a higher TNF/IL-10 ratio in HnpM-treated BMDCs **(Fig. 1D)**, consistent with a tolDC phenotype and published reports describing variable effects of helminth products on TLR signaling(5). Full CD4^+^ T cell activation requires TCR engagement with peptide-MHC and co-stimulatory signaling, yet tolerogenic DCs (tolDCs) express low levels of these molecules(6). HnpM-treated BMDCs displayed reduced MHC-II, CD86, and CD40 expression **(Fig. 1E)**, and CD11c_+_ splenic DCs from mice treated *in vivo* with HnpM had similar decreased expression of these markers **(Fig. 1F)**, confirming induction of a tolerogenic phenotype both *in vitro* and *in vivo*.

**Figure 1.**
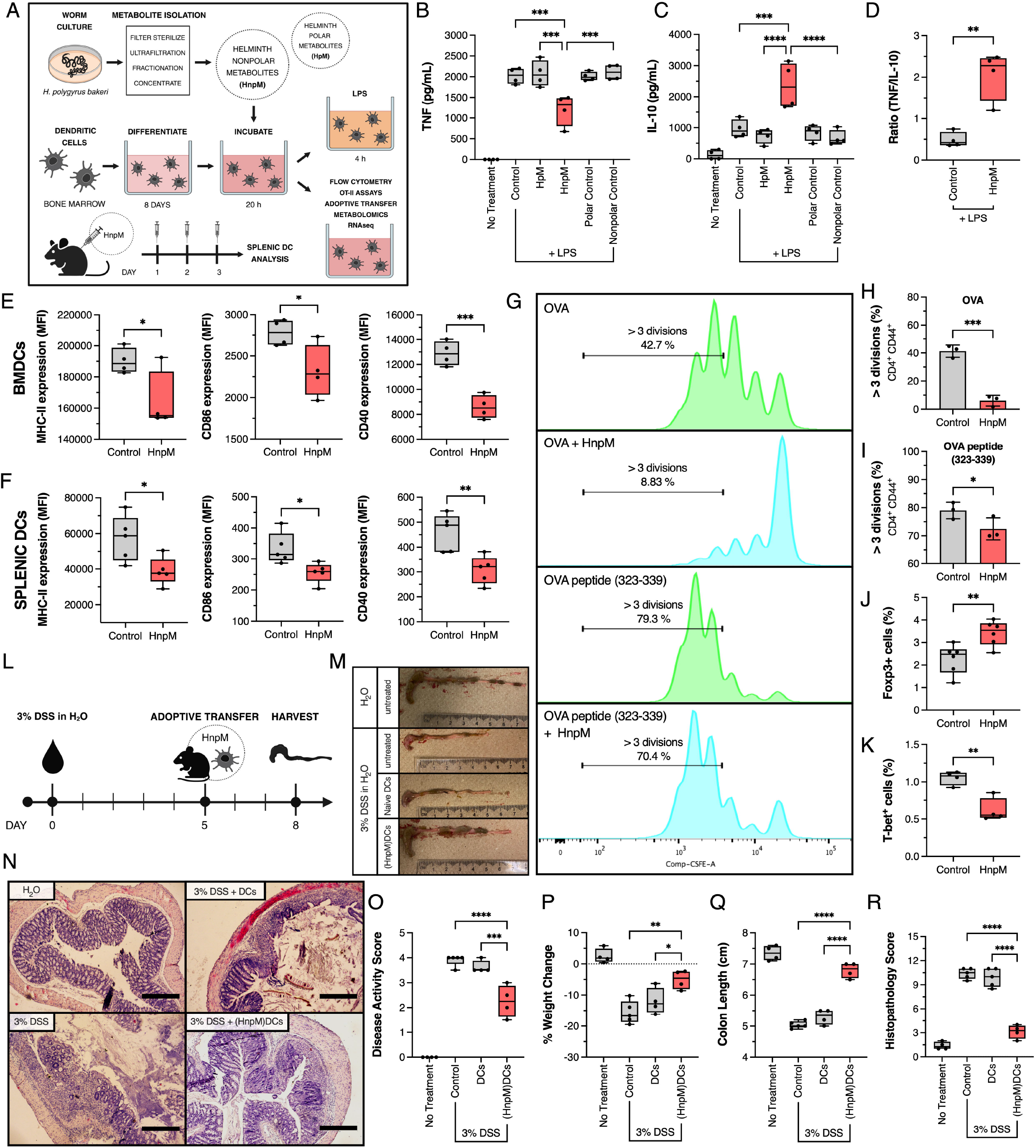
Helminth-derived nonpolar metabolites (HnpM) induce a tolerogenic phenotype in DCs. **(A)** Experimental design. BMDCs were treated with HpM or HnpM and challenged with LPS. ELISA quantifying **(B)** TNF, **(C)** IL-10, and **(D)** TNF/IL-10 ratio (n=4). Flow cytometry analysis of MHC-II, CD86, and CD40 expression from **(E)** HnpM-treated BMDCs and **(F)** CD11c_+_ splenic DCs isolated from mice treated with HnpM (n=4). HnpM-treated BMDCs inhibit antigen-specific CD4_+_ T cell proliferation. **(G)** Mean fluorescence intensity of CSFE dye dilution in gated CD4_+_CD44_+_ T cells and percentage of cells undergoing > 3 cell divisions (n=3). **(H, I)** Percentage of CD4_+_CD44_+_ T cells undergoing > 3 cell divisions. CD4_+_ T cells co-cultured with HnpM-treated BMDCs polarized towards a Treg phenotype. Flow cytometry analysis of **(J)** Foxp3_+_ expression and **(K)** T-bet+ cells (n=6). Adoptive transfer of HnpM-treated DCs ((HnpM)DCs) alleviate DSS-induced intestinal inflammation. **(L)** Mice were given 3% DSS in drinking water for 5 days followed by a one-time i.p. adoptive transfer of either 1 x 10_6_ naïve BMDCs or (HnpM)DCs. Quantification of **(O)** disease activity score, **(P)** percent weight change, **(M, Q)** colon length, and **(N, R)** histopathology scores, scale bar = 400 μm (n=4-5 mice). Data represented as boxplots and analyzed by one-way ANOVA with Holm-Sidak multiple comparisons **(B-C, O-R)** or unpaired *t* test **(D-F, H-K)**: ****p<0.0001, ***p<0.001, **p<0.01, *p<0.05.

To interrogate tolerogenic function *in vitro*, HnpM-treated BMDCs were pulsed with OVA or OVA peptide and co-cultured with CFSE-labeled OT-II CD4^+^ T cells. HnpM-treated DCs suppressed antigen-specific T cell proliferation **(Fig. 1G-I)**, promoted increased Foxp3^+^ Treg polarization **(Fig. 1J)**, and decreased T-bet+ Th1 polarization **(Fig. 1K)**. We assessed DC tolerogenic function *in vivo* by using a dextran sodium sulfate (DSS) model of intestinal inflammation in which B6 mice were given 3% DSS in the drinking water for 5 days, and HnpM-treated BMDCs [(HnpM)DCs] were adoptively transferred via i.p. injection on day 5 **(Fig. 1L)**. Mice that received (HnpM)DCs showed reduced colon shortening **(Fig. 1M, Q)**, improved histopathological scores **(Fig. 1N, R)** and disease activity scores **(Fig. 1O)**, and attenuated weight loss **(Fig. 1P)**, compared to DSS-treated mice receiving mock extraction-treated BMDCs. Although the immunosuppressive therapeutic repertoire has diversified for IBD, there is currently no approved therapy that effectively restores immune tolerance to alleviate intestinal inflammation(7). Our study reveals bioactive and immunomodulatory small molecules from helminths that are amenable to high throughput screening assays and further molecular characterization. We have previously reported that adoptively transferred BMDCs(8) or macrophages(9) migrate to lymphoid organs and the gastrointestinal tract, where they interact with T cells and alleviate intestinal inflammation. Others have demonstrated that tolDC adoptive transfer inhibits DNBS-induced colitis in immunocompetent but not in *Rag*^-/-^ mice(4), demonstrating that tolDC-T cell interactions are essential for inflammatory disease protection.

Once we established the functional activity of (HnpM)DCs, we determined their transcriptional signature, as it has been shown that immunogenic DCs and tolDCs express distinct transcriptional profiles(10). We compared HnpM-treated and mock extracted metabolite-treated BMDCs at 4 and 20h using RNAseq. Differential gene expression (DEG) analysis of HnpM-treated BMDCs against mock-treated BMDCs showed that at 4h there were no changes in gene expression **(Fig. 2A, C)**, but by 20h there were transcriptional changes in critical genes for DC function **(Fig. 2B, C)**. These genes may be investigated in the future as new targets of tolerance induction. Pathway enrichment analysis revealed features associated with immunity, including antigen processing and presentation, regulation of innate immune response, IL-10 production, and tolerance induction **(Fig. 2D)**, confirming the tolerogenic phenotype we observed. GSEA analysis revealed that DEGs identified in HnpM-treated BMDCs shared transcriptional similarities with genes previously identified to be upregulated in dexamethasone- and vitamin D3-induced tolDCs(11) **(Fig. 2E)**.

**Figure 2.**
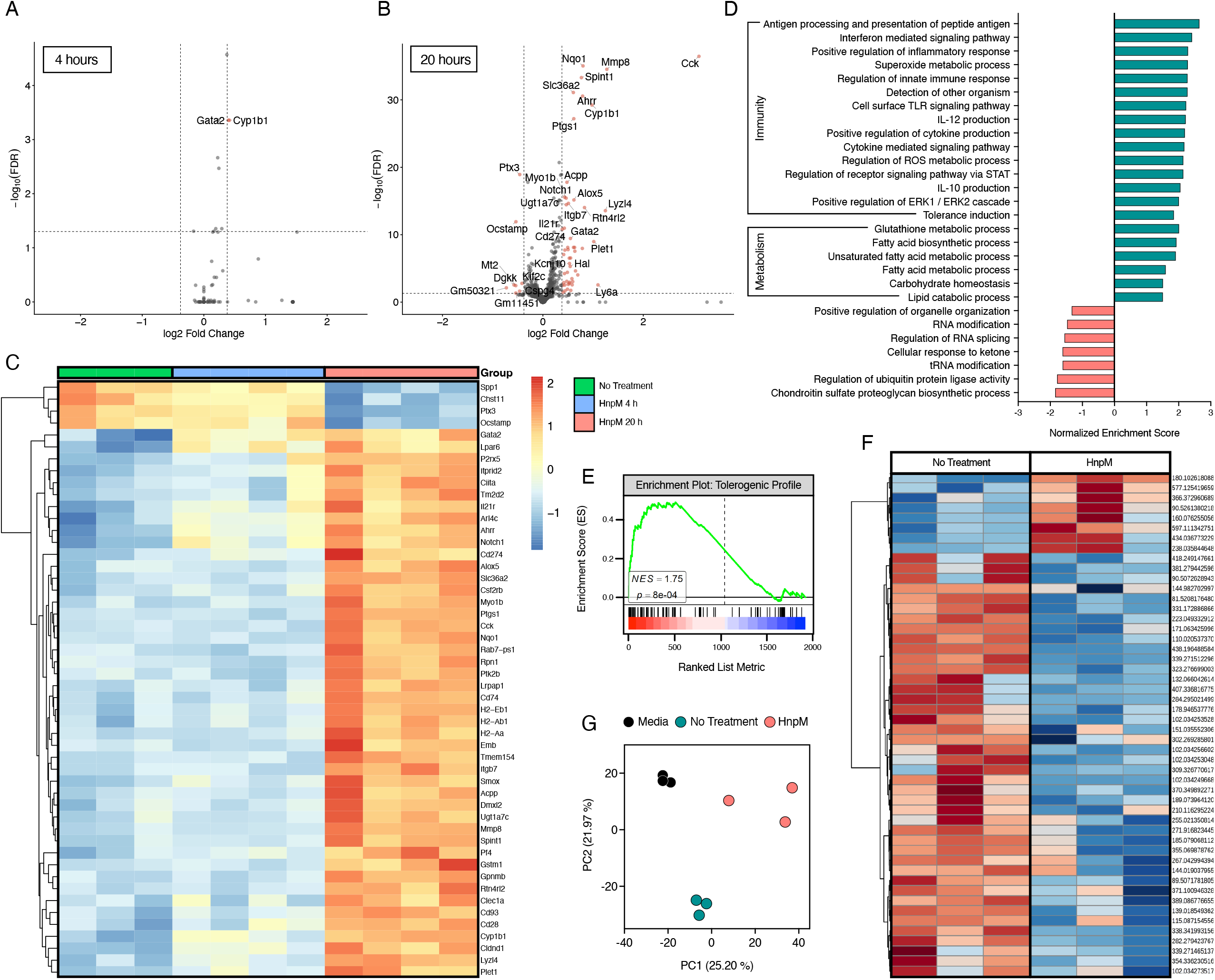
HnpM induces transcriptional and metabolic changes in BMDCs. BMDCs were incubated with HnpM for 4 and 20h and RNAseq and metabolomics were performed (n=4). Transcriptional profiling showing **(A, B)** volcano plots of DEGs at 4 and 20h, respectively, **(C)** heatmap of top 50 DEGs at 4 and 20h, and **(D)** Gene Set Enrichment Analysis of HnpM-treated BMDCs at 20h. **(E)** GSEA enrichment plot of tolerogenic gene set member positions across the ranked DESeq2 Wald statistic in the 20h HnpM-treated BMDCs. Untargeted global metabolomics revealed metabolite changes in HnpM-treated BMDCs as shown by a **(F)** heatmap of top 50 statistically enriched metabolite peak intensities at 4h, and **(G)** PCA plot of metabolites at 4h.

In addition to immunologically enriched pathways, we observed an enrichment of metabolic pathways. Metabolic reprogramming of immune cells is closely intertwined with inflammatory and tolerogenic responses(12). Thus, we performed untargeted global metabolomics on supernatants from HnpM- or mock-treated BMDCs at 4 and 20h. The majority of mammalian metabolites have uncharacterized biochemical structures and have bioactive functions that are not yet annotated. Thus, we comprehensively profiled the metabolite masses from HnpM-treated DCs to find potentially undescribed metabolite biomarkers of tolDCs. Using this unbiased approach, we observed that HnpM-treated BMDCs contained several unique metabolite peaks corresponding to a hierarchy of metabolic changes, in which we selected peaks using a 2-fold change cut-off against mock treated samples. Principal component analysis (PCA) indicated that BMDCs treated with HnpM for 4h formed distinct clusters from mock-treated or media control groups **(Fig. 2G)**. Metabolite analysis at 20h treatment did not reveal differences, suggesting that acute metabolic changes precede the transcriptional changes observed at 20h. At 4h, we observed a downregulated clustering of metabolites, as shown in the heatmap of the top 50 most enriched metabolites **(Fig. 2F)**. These observations support the conclusion that the early and rapid metabolic shift induced in HnpM-treated BMDCs precede gene expression changes associated with DC polarization towards a tolDC phenotype.

TolDCs promote immune homeostasis and tolerance by influencing CD4_+_ Th cell responses by numerous mechanisms. Recent studies indicate that different functional classes of DCs are marked by distinct phenotypic, molecular, and transcriptional signatures, which dovetail into diverse immunological functions within DC subsets(13). Collectively, our findings demonstrate that DC maturation and function are directly influenced by exposure to HnpM produced by adult stage Hpb. Helminth-derived metabolites that promote tolDCs likely evolved as immune evasion strategies enabling chronic infection. Our findings establish a foundation to define novel tolerance pathways and identify helminth small molecules and targets that could be harnessed to develop new therapies for inflammatory diseases.

## MATERIALS AND METHODS

### Mice

Male BALB/c mice, 8-10 weeks old, and female C57Bl/6 (B6) mice, 6-8 weeks old, obtained from Charles River Laboratories (St. Constant, QC, Canada) and B6 OT-II/Rag^-/-^ mice, 6-8 weeks old, generously provided by Dr. Judith Mandl (McGill University), were housed in a controlled environment, in a pathogen-free animal facility at McGill University, with a 12 h light-dark cycle at 23°C and 30 to 70% humidity, fed rodent chow *ad libitum*, and cared for by the Animal Research Department. All experiments were approved and conducted in accordance with the guidelines of the Canadian Council on Animal Care (Protocol # 2018-8066).

### *Heligmosomoides polygyrus bakeri* (Hpb)-derived small molecules

Helminth-derived excretory/secretory products were obtained as previously described(14). BALB/c mice were infected with 400-500 infective larvae (L3) of Hpb for 21 days. Adult worms were collected from the small intestine and cultured in RPMI 1640 medium containing 2% w/v glucose, 80 μg/ml gentamicin, 100 U/ml penicillin/streptomycin, and 20 μg/ml polymyxin B for 40 h at 37°C, 5% CO_2_. The supernatant was collected, centrifuged at 8000*g* for 10 min at 4°C, and passed through a 0.2 μm filter. Small molecules were separated from proteins and exosomes using a 3 kDa MWCO centrifugation unit (UFC910024, Millipore) and centrifuged at 4000*g* for 40 min at 4°C. Hpb-derived small molecules were further separated into polar (HpM) and nonpolar (HnpM) fractions by chromatography using a C_18_ column. The flow-through containing polar metabolites from the C_18_ column was further subjected to a universal polymeric reversed-phase sorbent containing column (WAT106202, Waters), and the polar metabolites were eluted using acetonitrile at 100% v/v. The stock fractions of HpM and HnpM were concentrated using a speed vacuum concentrator, resuspended in ultrapure water, and stored at -20°C.

### Dendritic Cells

Bone marrow-derived dendritic cells (BMDCs) and splenic dendritic cells were obtained from B6 mice. For BMDCs, bone marrow cell suspensions were cultured and differentiated in 10 cm non-tissue culture coated petri dishes in complete RPMI 1640 medium supplemented with 10% fetal bovine serum (FBS), 1% 100X penicillin/streptomycin, 1 mM sodium pyruvate, and 20 ng/ml mouse recombinant GM-CSF (78206, StemCell Technologies) for 8 days at 37°C, 5% CO_2_. On days 3 and 6, BMDCs were given an additional 10 mL and 5 mL, respectively, of complete RPMI medium containing 20 and 40 ng/ml GM-CSF. On day 8, the floating cells were collected and used for all experiments with flow cytometry confirming the purity of BMDCs as >90% CD11c_+_. BMDCs (1 x 10_5_ cells) were seeded into 96-well plates and treated with either 50 μg/mL HnpM, 50 μg/mL HpM, or 50 μg/mL polar and nonpolar mock extractions in complete RPMI 1640 media containing 20 ng/ml GM-CSF and cultured for 20 h at 37°C, 5% CO_2_. The BMDCs were rinsed and challenged with 100 ng/mL LPS (L2630, Millipore-Sigma) for 4 h. For splenic DC experiments, B6 mice were i.p. treated once per day for 3 days with either 50 μg/mL HnpM, 50 μg/mL HpM, or 50 μg/mL polar and nonpolar mock extractions. Spleens were harvested, minced, passed through a 40 μm cell strainer, and red blood cells were lysed in ACK buffer (1049201, ThermoFisher). The cells were resuspended in FACS buffer for flow cytometry staining and analysis.

### ELISA

Cytokine analysis on BMDC culture supernatants was performed using TNF and IL-10 ELISA DuoSets (DY410, DY417, R&D Systems) according to the manufacturer’s recommendation.

### Immunophenotyping

Cultured BMDCs and splenic DCs were incubated in F_c_ block and labeled with the following antibodies from BioLegend for 30 min at 4°C: anti-CD45-PeCy7, -CD11c-FITC), -MHC-II (I-A/I-E)-BV421, -CD40-PE, and -CD86-PerCP mAbs. To quantify CD4_+_Foxp3_+_ cells, spleen cells were stained with anti-CD4-FITC mAb (eBioscience), fixed, and permeabilized using the Foxp3 staining kit (Invitrogen), and stained with anti-Foxp3-Alexa Fluor 488 mAb (5523, eBioscience) for 30 min at 4°C. Cell events were acquired using an Attune NxT flow cytometer (ThermoFisher Scientific) and analyzed using FlowJo software v.10.2 (Tree Star).

### OVA-specific CD4^+^ T cell proliferation

To evaluate OVA-specific CD4_+_ T cell proliferation, spleens were harvested from naïve OT-II TCR-transgenic mice, and single cell suspensions were prepared as described above. CD4_+_ T cells were isolated by negative selection using an EasySep Mouse CD4_+_ T cell Isolation Kit (19852, StemCell Technologies) according to the manufacturer’s recommendation. BMDCs were either treated with 50 μg/ml HnpM or mock treated for 24 h prior to co-culture with OT-II CD4_+_ T cells. During the final 6 h of culture (18 h), BMDCs were pulsed with either 250 μg/mL OVA (Sigma) or 10 μg/mL OVA peptide 323-339 (T78000, Sigma) at 37°C, 5% CO_2_. At 24 h, BMDCs were co-cultured with purified, CFSE-labelled (34554, eBioscience) OT-II CD4_+_ T cells at a final T cell to BMDC ratio of 2:1 for 72 h. Cells were collected and labelled with anti-CD4-FITC and anti-CD44-APC mAbs (BD Biosciences). Flow cytometry was performed to determine CFSE dilution in gated CD4_+_CD44_+_ cells (LSR Fortessa; BD Biosciences), and data was analyzed using FlowJo software.

### DSS Colitis

Female B6 mice 6-8 weeks old were provided with 3% w/v DSS (MP Biomedicals) in the drinking water *ad libitum* for 5 days. On day 5, the drinking water was replaced with normal drinking water and mice were injected i.p. with 1 x 10_6_ BMDCs (PBS vehicle control), naïve DCs, or HnpM-treated DCs ((HnpM)DC)). Mouse weight was monitored and recorded daily. On day 8, necropsies were performed and colon lengths, percent weight changes, histopathology scoring, and disease activity scores (DAS) based on a scale of 0-5 were determined based on body weight, colon length, health of the animal, rectal bleeding, and macroscopic appearance of the colon(15).

### Histology

Mid-distal colon sections were fixed in 10% neutral buffered formalin, embedded in paraffin wax, and sectioned to 4 μm thickness. Sections were de-paraffinized in two changes of xylene and rehydrating through graded ethanol solutions (100%, 95%, 70% EtOH) and distilled H_2_O. Sections were stained in Gills II hematoxylin (50-255-2441, Leica Biosystems) for 5 minutes, rinsed in lukewarm tap water for 5 min, counterstained in eosin (3801600, Leica Biosystems) for 1 min, followed by 1 min in distilled H_2_O. The sections were dehydrated through reverse graded ethanol solutions (70%, 95%, 100% EtOH) and cleared in two changes of xylene before mounting with Permount (SP15, Fisher Scientific). Brightfield microscopy on mid-distal histological sections was performed using a 100X objective on an EVOS XL Cell Imaging System (ThermoFisher Scientific) in which 5 mid-distal colon sections per mouse per group were blindly assessed using a validated 12-point scoring system.

### Metabolomics and Transcriptomics Experiments

BMDCs were cultured as previously described and treated with either HnpM or vehicle control for 4 and 20 h and collected for matched transcriptomics and metabolomics analysis. For transcriptomics, RNA from HnpM-treated and vehicle control treated BMDCs was extracted in TRIzol (15596026, Invitrogen) and chloroform and precipitated with isopropanol and extracted using PureLink™ RNA Micro Scale Kit (12183016, ThermoFisher). For metabolomics, supernatants and cell-free media controls were extracted using ice-cold HPLC-grade methanol for 30 min on ice, centrifuged at 10,000*g* for 10 min at 4°C, and stored at -80°C until analysis.

### RNA sequencing

RNA sequencing of poly-A enriched RNAs was performed at Centre d’expertise et de services Génome Québec, as previously described(16). Reads were aligned to *Mus musculus* reference genome GRCm38.p6 using STAR aligner and gene-level counts were generating using featureCounts(17). Differential gene expression analysis was conducted in R using the DESeq2 package(18). Genes with fewer than 10 raw counts were filtered prior to analysis, p-values were adjusted for multiple testing using the Benjamini-Hochberg method, and genes were considered significantly differentially expressed at an adjusted p-value < 0.05 and fold-change greater than 1.5. Heatmaps were constructed using VST-transformed data and row-wise z-score scaling. Gene Set Enrichment Analysis (GSEA)(19) was performed using the fgsea package in R using the Wald statistic as the ranked metric. Gene sets were obtained from MSigDB.

### Metabolomics

Nonpolar metabolite profiling was performed at the UVic-Genome BC Proteomics Centre by LC-MS/MS analysis of the deproteinated conditioned media by injecting 3 mL of sample onto a Dionex UHPLC system equipped with an Agilent Eclipse C_18_ (2.1 ⨯ 15 mm, 1.8 mm) column incubated at 45°C. Metabolites were resolved with a 30 min linear running 0-80% using the buffer system 0.05% formic acid and 0.05% formic acid in acetonitrile at a flow rate of 300 mL/min. The column effluent was introduced by electrospray ionization onto a Velos LTQ Orbitrap Analyzer (Thermo Scientific) using a spray voltage of 3.6 kV, a source heater temperature of 350°C, and a sheath gas flow of 40 l/min. Survey scans were performed using the Orbitrap mass spectrometer and the 10 most intense ions were selected for fragmentation using a 30-40 V-stepped collision-induced dissociation energy. Fragmentation products were analyzed in the linear ion trap mass spectrometer. Fragmentation was used to perform an online search using the XCMS database (https://xcmsonline.scripps.edu) to identify possible metabolites. Following peak profiling, 2-fold change enriched peaks (compared to control samples, i.e., samples with media only) were selected to perform statistical and functional interpretation using MetaboAnalyst 6.0 (https://www.metaboanalyst.ca/). Data were analyzed using interquartile range (IQR) and normalized by log_10_ transformation. Principal component analysis (PCA) and heatmap (with Euclidean distance measure and Ward clustering algorithm with clustered group samples) were performed. Peak annotations were further analyzed using Mummichog algorithm and KEGG database for functional interpretation.

### Statistical analysis

Statistical analyses of metabolomic and transcriptomic data are described in the corresponding method sections. All other data are presented as boxplots indicating the median (center line), upper and lower box bounds (IQR = first and third quartiles), and whiskers (minimum and maximum values), with individual data points superimposed onto the boxplot. Normality of datasets were tested using the Kolmogorov-Smirnoff test. Data that were normally distributed were analyzed by one-way ANOVA with appropriate post-hoc tests. Unpaired data were compared using two-tailed unpaired Student’s *t* test. Statistical analysis was performed using GraphPad Prism v.11.0.0, and significance was assumed for p < 0.05.

## DATA AVAILABILITY

Metabolite LC-MS data was deposited to Metabolomics Workbench under accession number ST002230. Transcriptomics data (RNAseq) was deposited to the NCBI Sequence Read Archive (SRA) under accession number PRJNA856720.

## AUTHOR CONTRIBUTIONS

Conceptualization, A.N.M., N.L.M., A.J., M.M.S., F.L.; Data Curation, A.N.M., N.L.M., J.A.S., T.S., G.F., I.L.K., I.C., A.J., F.L.; Data Analysis, A.N.M., N.L.M., J.A.S., T.S., G.F., I.L.K., I.C., A.J., F.L.; Funding Acquisition, M.M.S., T.H., F.L.; Investigation, A.N.M., N.L.M., J.A.S., G.F., I.L.K.; Methodology, A.N.M., N.L.M., J.A.S., E.S., A.G.M., T.S., G.F., I.L.K., I.C., A.J., M.M.S., F.L.; Project Administration, F.L.; Resources, I.L.K., M.M.S., T.H., F.L.; Software, I.C.; Supervision, M.M.S, F.L.; Validation, F.L.; Visualization, A.N.M., J.A.S.; Writing - Original Draft, A.N.M., N.L.M., F.L.; Writing - Review & Editing, A.N.M., N.L.M., J.A.S., E.S., A.G.M, T.S., G.F., I.L.K., I.C., A.J., M.M.S., T.H., F.L.

## COMPETING INTERESTS

The authors declare no competing interests.

## ACKNOWLEDGEMENTS

This study was supported by the Natural Science and Engineering Research Council of Canada (NSERC) (grant numbers RGPIN/04459-2019 to F.L. and RGPIN/1327730 to M.M.S.), Canadian Institutes Health Research (CIHR) (grant number MOP130369) to M.M.S., and Fonds de Recherche du Québec – Nature et Technologies (FRQ-NT) (grant number 253419) to M.M.S. and F.L. A.N.M was supported by NIH T32 (2T32CA009370-40).

## Notes

https://www.metabolomicsworkbench.org/data/DRCCMetadata.php?Mode=Project&ProjectID=PR001420

### Competing Interest Statement

The authors have declared no competing interest.

### Summary of Updates

New analyses were performed on RNAseq datasets. Figures have been condensed and updated. The manuscript has been rewritten to reflect updated analyses and figures. The author list has been amended to add contributing authors. Author affiliations updated.

## REFERENCES

1. G. G. Kaplan, IBD: Global variations in environmental risk factors for IBD. Nat Rev Gastroenterol Hepatol 11, 708–709 (2014).

2. T. Setiawan et al., Heligmosomoides polygyrus promotes regulatory T-cell cytokine production in the murine normal distal intestine. Infect Immun 75, 4655–4663 (2007).

3. M. Arora et al., Simvastatin promotes Th2-type responses through the induction of the chitinase family member Ym1 in dendritic cells. Proc Natl Acad Sci U S A 103, 7777–7782 (2006).

4. C. E. Matisz et al., Adoptive transfer of helminth antigen-pulsed dendritic cells protects against the development of experimental colitis in mice. Eur J Immunol 45, 3126–3139 (2015).

5. C. M. Kane et al., Helminth antigens modulate TLR-initiated dendritic cell activation. J Immunol 173, 7454–7461 (2004).

6. V. K. Raker, M. P. Domogalla, K. Steinbrink, Tolerogenic Dendritic Cells for Regulatory T Cell Induction in Man. Front Immunol 6, 569 (2015).

7. S. Danese, G. Fiorino, L. Peyrin-Biroulet, Positioning Therapies in Ulcerative Colitis. Clin Gastroenterol Hepatol 18, 1280-1290.e1281 (2020).

8. C. E. Matisz et al., Helminth Antigen-Conditioned Dendritic Cells Generate Anti-Inflammatory Cd4 T Cells Independent of Antigen Presentation via Major Histocompatibility Complex Class II. Am J Pathol 188, 2589–2604 (2018).

9. J. L. Reyes et al., Macrophages treated with antigen from the tapeworm Hymenolepis diminuta condition CD25(+) T cells to suppress colitis. Faseb j 33, 5676–5689 (2019).

10. B. Vander Lugt et al., Transcriptional determinants of tolerogenic and immunogenic states during dendritic cell maturation. J Cell Biol 216, 779–792 (2017).

11. J. Navarro-Barriuso et al., Comparative transcriptomic profile of tolerogenic dendritic cells differentiated with vitamin D3, dexamethasone and rapamycin. Sci Rep 8, 14985 (2018).

12. R. Stienstra, R. T. Netea-Maier, N. P. Riksen, L. A. B. Joosten, M. G. Netea, Specific and Complex Reprogramming of Cellular Metabolism in Myeloid Cells during Innate Immune Responses. Cell Metab 26, 142–156 (2017).

13. A. C. Villani et al., Single-cell RNA-seq reveals new types of human blood dendritic cells, monocytes, and progenitors. Science 356 (2017).

14. R. M. Valanparambil et al., Production and analysis of immunomodulatory excretory-secretory products from the mouse gastrointestinal nematode Heligmosomoides polygyrus bakeri. Nat Protoc 9, 2740–2754 (2014).

15. F. Lopes et al., The Src kinase Fyn is protective in acute chemical-induced colitis and promotes recovery from disease. J Leukoc Biol 97, 1089–1099 (2015).

16. A. Rodríguez-García, A. Sola-Landa, R. Pérez-Redondo, Coupled Transcriptomics for Differential Expression Analysis and Determination of Transcription Start Sites: Design and Bioinformatics. Methods Mol Biol 2296, 263–278 (2021).

17. A. Dobin et al., STAR: ultrafast universal RNA-seq aligner. Bioinformatics 29, 15–21 (2013).

18. M. I. Love, W. Huber, S. Anders, Moderated estimation of fold change and dispersion for RNA-seq data with DESeq2. Genome Biol 15, 550 (2014).

19. A. Subramanian et al., Gene set enrichment analysis: a knowledge-based approach for interpreting genome-wide expression profiles. Proc Natl Acad Sci U S A 102, 15545–15550 (2005).

